# GreenSloth: a curated database and executable platform for mechanistic photosynthesis models

**DOI:** 10.64898/2026.07.22.740007

**Authors:** Elouen Corvest, Marvin van Aalst, Tim Nies, Quang Huy Nguyen, Josha Ebeling, Maja Strauch, El Hadji Malick Cisse, Tanvir Hassan, Anna Matuszyńska

## Abstract

Mechanistic models of photosynthesis have expanded substantially over the past decades, covering processes from light reactions to carbon fixation. However, these models remain fragmented across the literature, inconsistently implemented, and difficult to reproduce or reuse, limiting their adoption beyond the research group that developed them.

Here, we present GreenSloth, a freely accessible web-based database of 22 published mechanistic photosynthesis models, reimplemented as standardized, executable Python objects within MxlPy, an open-source framework for mechanistic biological modeling. Although the database is primarily designed for dynamic mechanistic models formulated as ordinary differential equations, the current implementation also includes the field’s most widely cited steady-state mechanistic model and its variants.

GreenSloth provides a structured environment for model discovery, comparison, and reuse, addressing reproducibility challenges in the field and enabling integration into emerging hybrid modeling approaches. It is also interactive: each model runs directly in the browser, with no installation, environment setup, or programming required. The resource is openly accessible and designed for long-term community maintenance, hoping to position itself as foundational infrastructure for the photosynthesis modeling community.

**Database URL:** https://greensloth.rwth-aachen.de/

## 1 Introduction

Photosynthesis is fundamental to global carbon fluxes and agricultural productivity. Consequently, extensive research has been devoted to understanding its underlying mechanisms through both experimental and theoretical approaches [1, 2]. Dynamic models, formulated as systems of ordinary differential equations (ODEs), have been particularly instrumental in advancing our understanding of photosynthetic regulation, ranging from compact representations of a single mechanism to large networks coupling electron transport, photoprotection, and carbon fixation [3]. Recently, the aim of photosynthesis modeling has increasingly shifted from photosynthesis under a controlled environment towards the response under natural environments with multiple fluctuating factors [4, 5, 6]. Under natural and agricultural conditions, plants experience changes in light, temperature, CO_2_, and water availability, while key physiological processes, including ribulose-1,5-bisphosphate carboxylase/oxygenase (RuBisCO) activation, stomatal conductance, non-photochemical quenching (NPQ), and metabolic regulation, respond on different timescales [7]. The complex responses of photosynthesis to multiple environmental factors require a systematic framework to disentangle the effects of individual factors and their interactions on plant performance.

Despite decades of accumulated models that span widely different scopes, structures, and applications (partially reviewed in [2, 3, 8, 9, 10]), no dedicated resource exists to collect them, nor to allow for easy comparison or standardized usage. Implementations remain scattered, inconsistently reported, and difficult to reproduce, and prior platforms have struggled with long-term maintenance (e-Photosynthesis [11]) or adoption in this field specifically (BioModels [12]). The dispersion of models is costly for three distinct user groups. Computational biologists developing mechanistic models of photosynthetic processes often must reimplement models from scratch for each new study, rather than building on validated, reusable code. Researchers from the scientific machine learning community developing hybrid or physics-informed approaches have no single place to access validated first-principles models to merge with data-driven techniques. Finally, experimental collaborators, lacking programming expertise, cannot easily identify which existing model matches their system or hypothesis before designing an experiment. Currently, no dedicated resource addresses all three needs simultaneously.

For this purpose, we developed GreenSloth, a curated database of executable Python-based photosynthesis models. To ensure reproducibility, GreenSloth is built on MxlPy [13], an open-source Python framework for implementing, simulating, and analyzing mechanistic biological models. Critically, GreenSloth does not merely archive model code: each model can be run directly in the browser via mxl-web [14], without installation, environment setup, or programming required. This removes a major practical barrier to model reuse, allowing experimental collaborators and ML practitioners alike to interact with a validated mechanistic model within seconds of finding it, and is consistent with the broader case for web-based, in-browser scientific computing we outlined in [14]. The database classifies models using process-specific tags and provides structured information on their equations, parameters, and implementations, allowing researchers to compare models and identify the one best suited to their research question.

## 2 Materials and Methods

### 2.1 Model selection

Mechanistic and dynamic photosynthesis models were initially identified from published literature. The reviews of Stirbet *et al*. [2, 3], *von Caemmerer [8], Sukhova et al*. [9], and Jablonsky *et al*. [10] provided a comprehensive view of the main approaches and open questions for the photosynthesis modeling community, guiding the curation process. These models were shortlisted for GreenSloth based on relevance, citation frequency, and coverage of key photosynthetic processes. Models passing an initial relevance test were then evaluated against five formal eligibility criteria (see Fig. 1). They (i) described photosynthetic processes using a mechanistic mathematical formulation, (ii) were formulated as dynamic systems of ODEs, (iii) were published in peer-reviewed literature, (iv) reported sufficient equations and parameter values to permit reconstruction, and (v) could be validated against published simulations. Criterion (ii) was relaxed for one exception: the FvCB model [15] and its expansions, as the field’s most cited steady-state framework, were included despite not meeting the dynamic-ODE requirement, given their historical influence. The final model list was established to cover central aspects of photosynthesis: the light reactions, electron transport chains, the carbon fixation process, the chloroplast proton motive force (*pmf* ), photorespiration, and the photoprotective NPQ processes.

**Figure 1:**
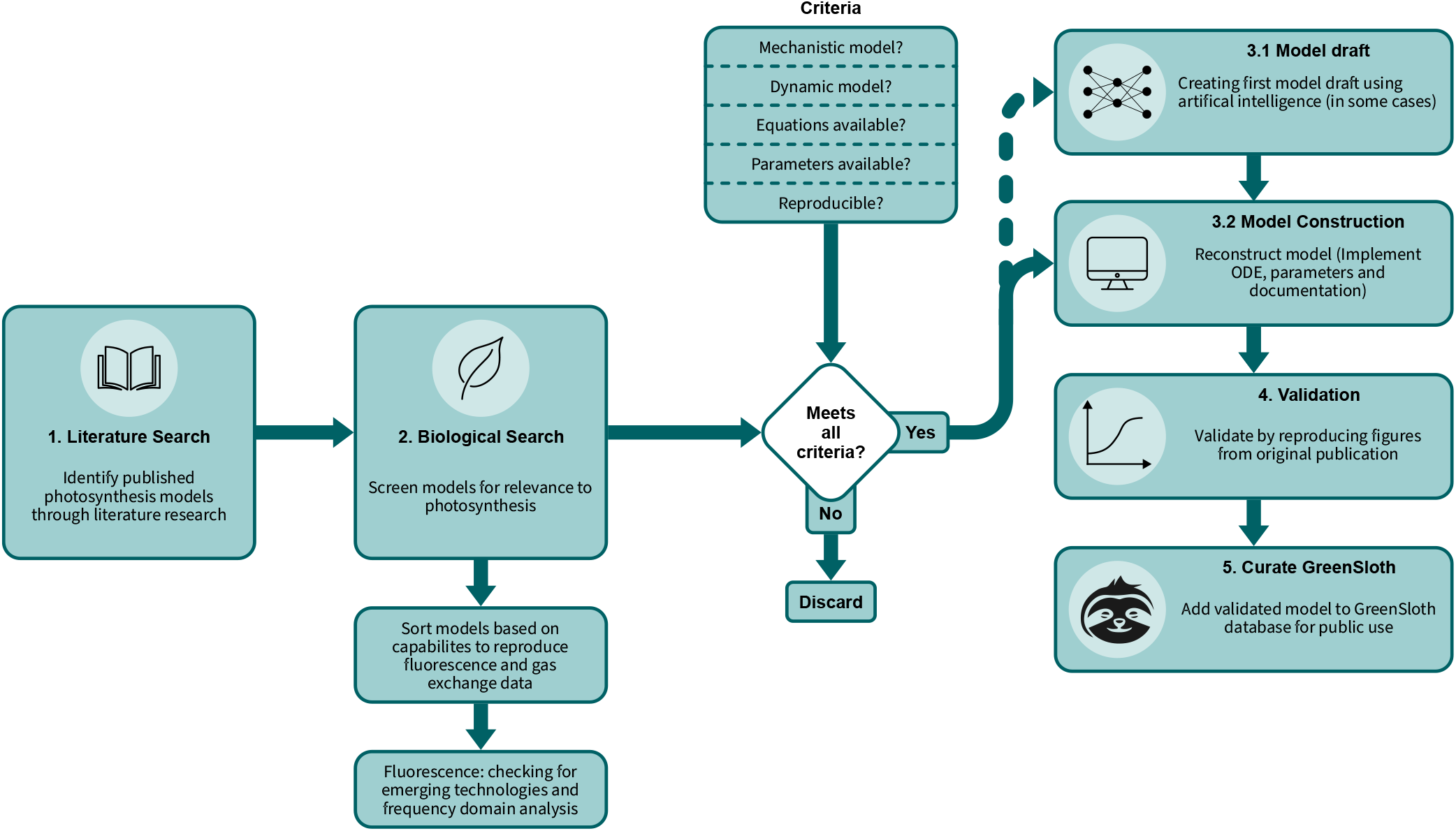
Workflow for selecting mechanistic and dynamic photosynthesis models for GreenSloth. Candidate models were first identified through a literature search and screened for biological relevance, with priority given to models capable of reproducing fluorescence and gas-exchange data. Models passing this screen were evaluated against five formal eligibility criteria (mechanistic formulation, dynamic ODE structure, peer-reviewed publication, sufficient reported equations and parameters, and validity) before reconstruction in MxlPy [13]. Each reimplementation was validated by recreating the corresponding figures from the original publication, with any discrepancies documented. Validated models were then curated into the GreenSloth database, with all relevant information made publicly available.

### 2.2 Model implementation

Each model was reimplemented in MxlPy [13] using the parameter values, units, and initial conditions reported in the original publication, obtained either from freely available model code or from the supplementary materials and main text. Where necessary, underlying assumptions not explicitly stated in the original publication were made explicit and documented in the model’s source code.

### 2.3 Model validation

Each reimplementation was validated against its original publication by recreating the corresponding figures and simulated trajectories, which served as the reference standard. A model was considered validated when its reimplementation visually reproduced these published results. Where discrepancies arose, typically due to missing or ambiguous parameters, undocumented units, or sign errors in the reported equations, these were resolved where the source of the error could be identified, and the correction was documented directly on the model’s page. Where a discrepancy could not be fully resolved, it was documented as a partial or qualitative reproduction rather than treated as a validation failure.

### 2.4 Platform implementation

GreenSloth is maintained in a single code repository and developed as a web-first interface. The platform has no backend server, which simplifies maintenance and allows deployment directly on GitHub Pages. The frontend is built with Svelte [16], a user interface framework that enables the creation of highly optimized websites. Svelte makes it easier to reuse layouts and styles, especially for pages that follow the same structure, such as each model page in the database.

Each model consists of a directory containing a Markdown file with the model summary, a TypeScript file with all the meta information of the model, another TypeScript file with a compatible version of the model, and some assets for the model page. With a consistent structure for the model directories, Svelte can automatically extract all the information needed to inject it into the template of the model’s web page.

The webpage of each model (see Fig. 2) includes information about the model authorship (original publication and reimplementation), model summary and scheme, a small live analysis, tables of important model aspects, and reproduced figures from the original publication for the validation of the implementation. The live analysis is performed using the Python package mxl-web [17].

**Figure 2:**
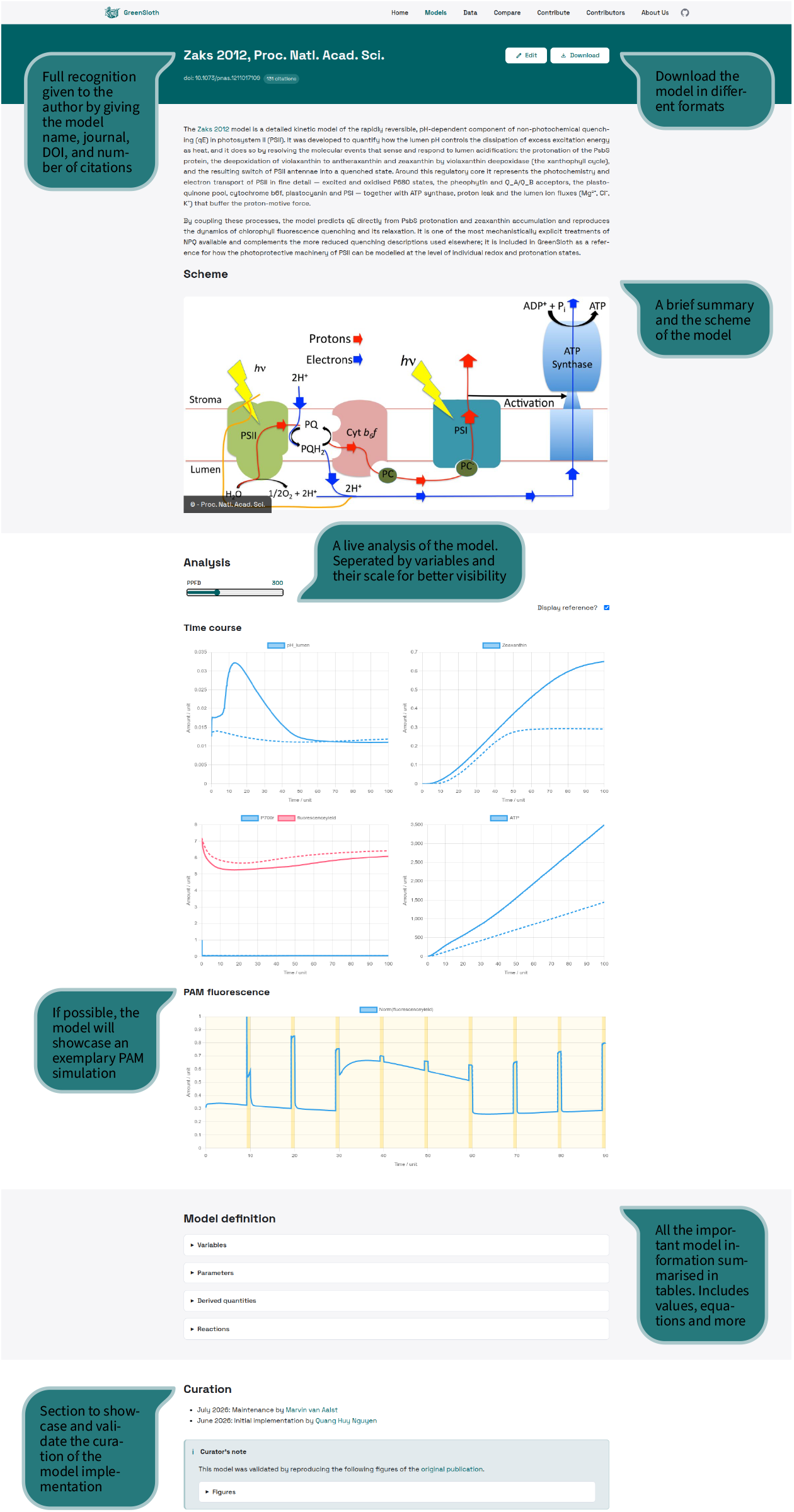
Overview of a model page on GreenSloth. The Zaks2012 model page [18] is shown as a representative example of the GreenSloth model-page layout, which follows the same template across all models in the database. Annotations highlight key features: recognition of model authorship, a direct model download, a summary and scheme, a live analysis, an exemplary pulse amplitude modulation (PAM) simulation, tables of key model information, and a curation section documenting the validation of the reimplementation. The full page is available at https://computational-biology-aachen.github.io/green-sloth/models/zaks2012.

### 2.5 Model search

GreenSloth classifies each model using a tag system spanning four categories: (i) the photosynthetic process the model description includes (e.g., electron transport, NPQ, carbon fixation), (ii) the model type (dynamic ODE-based, steady-state), (iii) the type of experimental data the model can simulate or reproduce (e.g., fluorescence, gas exchange), and (iv) information on organism for which the model was developed (e.g., generic C3 plant model, *Arabidopsis thaliana*, or *Chlamydomonas reinhardtii* ). Short descriptions of each data type are provided on the GreenSloth website to help users unfamiliar with a given experimental technique interpret this category. Furthermore, these tags are integrated into an interactive scheme of the photosynthetic apparatus, allowing users to filter models directly by selecting a specific process (Section 3.2).

### 2.6 Contribution

GreenSloth is designed as a continuously updated, community-maintained database. Contributed models must follow the same directory structure described in in Section 2.4, detailed in a contribution guide on the website. Experienced users can submit a pull request to the “green-sloth” GitHub repository, manually creating a correctly formatted model directory. Alternatively, contributors can use an on-site submission tool to input the required information directly; on completion, it redirects the contributor to the GitHub repository with a pre-filled issue for the new model. Because the website has no backend server (Section 2.4), both submission routes are finalized through the GitHub repository. Each submission, whether a pull request or an issue, is reviewed by the repository maintainers against the same eligibility and validation criteria applied to the initial model collection (Sections 2.1–2.3). Once any requested changes are made, the submission is accepted, and the model becomes visible on the website.

## 3 Results

### 3.1 Model integration and validation

The first public release of GreenSloth (v0.1, archived at 10.5281/zenodo.21372931) comprises 22 mechanistic and dynamic models of photosynthesis, successfully reimplemented and integrated into the platform. The scope of these models ranges from mechanistic models of fluorescence induction to broad-scale simulation of the interaction between the photosynthetic electron transport chain and the Calvin-Benson-Bassham cycle (complete list available in Supplementary Table 1). Of the 22 models in this release, 14 were validated directly, reproducing the original published trajectories without modification; the remaining 8 required correcting a missing or ambiguous parameter, an undocumented unit, or a sign error before validation succeeded. Nevertheless, all models in the GreenSloth database can be considered validated against their original publication; whether a given model is sufficiently validated against experimental data for a specific research question remains open and is left to the user to decide.

### 3.2 Model discovery and comparison

To facilitate access to GreenSloth, a user-friendly website has been created. Once a user enters GreenSloth, a landing page welcomes them, explaining the motivation behind the database and key aspects of the platform. From there, the user can navigate to the “Models” page, which lists all models currently in the database. To help with the search, different systems have been implemented, all described in Section 2.5. Once the search has been narrowed down, the grid view of model tiles shows those that match, where each tile displays the model name, the journal in which it was published, and its corresponding scheme. These tiles are clickable, forwarding the user to the respective model page. Once a model is chosen (see Fig. 2), the user is greeted by a banner that includes the publication’s DOI, a live citation count from the OpenCitations API, and a download button offering various file formats. The banner is an important part of each model page, as GreenSloth does not take ownership of the models; rather, it aims to support model authors in reaching new audiences. Below the banner, the user can find more detailed information about the model: a summary, the model scheme (if available), live analysis options, the model definitions, and validation of the reimplementation. The model definitions consist of tables for important aspects of a mechanistic model, such as variables and parameters, with mathematical symbols written in L^A^TEX, IDs, and initial values or equations. To facilitate comparisons between models, the GreenSloth website provides a comparison page where users can select two models for a side-by-side comparison of variables, parameters, and other implementation information, allowing users to obtain a quick overview of what each model entails without inspecting its source code.

#### Text Box 1

**Working example**

Consider a researcher investigating short-term photoprotective responses under fluctuating light who wants to identify an existing mechanistic model of ()o guide the design of a new experiment. Filtering GreenSloth by the “NPQ/photoprotection” process tag returns several candidates, including the Matuszyńska *et al*. (2016) model of the xanthophyll cycle and short-term light memory [19] and the Zaks *et al*. (2012) kinetic model of rapidly reversible quenching [18]. Selecting these two models in the comparison tool surfaces their shared focus on qE-type quenching alongside differences in mechanistic scope, allowing the user to judge which model’s assumptions best match their experimental system without inspecting either model’s source code directly.

### 3.3 Live simulation

Each model page includes a small live-in-web simulation powered by mxl-web [14], which allows running ODE models entirely client-side, with no installation required. While mxl-web lets users edit and create their own models, only a subset of the toolbox’s functionalities is available on GreenSloth, restricted to live simulations with a small number of adaptable parameters.

Parameter values can be changed using integrated sliders. Most model pages allow for changing the photosynthetic photon flux density (PPFD) density, but depending on the model, other parameters may be included in the live simulations. The model contributor chooses which plots should be displayed on the model page. At the time of writing, the available plot types are: a time-course simulation, a pulse amplitude modulation (PAM) simulation, and a two-parameter analysis for steady-state models. By default, the time-course simulation visualizes all available model variables, often grouped by magnitude of the simulated results (e.g., concentrations) for better visualization. The PAM simulation uses an exemplary saturating pulse protocol and displays the model’s simulated fluorescence signal. A working example: Having the Zaks2012 model [18], the user can run its live simulation directly on the model’s page. Using a slider to change the input to the PPFD allows direct observation of the changes in the model’s time-course simulation. Additionally, this model can simulate a PAM protocol, as shown here under a standard PAM protocol (see Fig. 2), providing an immediate, interactive sense of the model’s dynamic behavior before committing to a full local installation. For steady-state applications, an analogous workflow applies to the Farquhar, von Caemmerer, and Berry (FvCB) model [15]: filtering by “carbon fixation” and “steady-state” retrieves the FvCB model and its variants [8, 20], and the live two-parameter analysis lets a user visualize, for example, the effect of CO_2_ concentration and PPFD on net assimilation rate directly in the browser.

## 4 Discussion and conclusion

GreenSloth addresses a practical and persistent limitation in photosynthesis modeling: mechanistic models are widely available in the literature, but they are rarely available as standardized, executable, and directly comparable resources. By curating published ODE-based models into a common framework, GreenSloth lowers the barrier for model reuse, validation, and further model development, directly addressing the three user groups motivating this work: theoretical/computational biologists, researchers from the scientific machine learning community, and experimental collaborators.

For computational biologists, a central value of the database is that it makes model assumptions explicit and provides a validated, reusable starting point rather than requiring reimplementation from scratch. During curation, equations, parameters, state variables, units, initial conditions, and simulation protocols are reconstructed and documented. This process exposes ambiguities that are often hidden in conventional model publications, including missing parameter values, unclear units, incomplete initial conditions, or insufficiently described simulation settings. GreenSloth therefore functions not only as a repository but also as a reproducibility audit of the photosynthesis modeling literature. This process exposes ambiguities that are often hidden in conventional model publications, including missing parameter values, unclear units, incomplete initial conditions, or insufficiently described simulation settings. GreenSloth therefore functions not only as a repository but also as a reproducibility audit of the photosynthesis modeling literature. Because curated models are implemented within a common computational environment, users can also examine differences in biological scope, mathematical formulation, parametrization, and dynamic behaviour, which is particularly relevant for models of photosynthetic electron transport, photoprotection, proton motive force, and carbon fixation, where different modeling traditions often describe overlapping processes using incompatible assumptions or levels of abstraction. Beyond comparison, because all models share a common implementation structure within MxlPy, GreenSloth also lowers the barrier to composing models of different photosynthetic subprocesses into larger integrated systems. A user can, for example, couple a light-reaction model with a carbon-fixation model with substantially less manual reconciliation than combining two independently implemented codebases, since both follow the same structural conventions and can be understood and connected in a plug-and-play manner.

GreenSloth can also be used to support emerging hybrid modeling approaches applied by researchers from the scientific machine learning community. Hybrid approaches such as universal differential equations and physics-informed neural networks combine a mechanistic core with learned components, using the mechanistic structure to constrain predictions, preserve interpretability, and enable extrapolation beyond the training data [21]. Building such models requires a validated, executable mechanistic component to start from. Reconstructing one from a published equation set demands modeling expertise that most researchers in this community do not have, and even when models have been deposited as systems biology markup language (SBML) files, this is no guarantee of usability. BioModels’ own reproducibility audit found that only 51% of surveyed models could be directly reproduced from the information reported in their original publications, with the remainder requiring manual correction or, in over a third of cases, remaining irreproducible entirely [22]. GreenSloth removes this barrier by providing ready-to-use, validated mechanistic models directly in Python, positioning it as a practical entry point for physics-informed and hybrid photosynthesis modeling.

Last, but not least, for experimental collaborators without programming expertise, GreenSloth’s browser-based execution (Section 3.3) removes the barrier that has traditionally separated model users from model builders. Rather than requiring familiarity with a specific programming environment to inspect a model’s predicted behavior, a researcher can filter directly to a model matching their biological system, run its live simulation, and evaluate whether its assumptions fit their experimental hypothesis before committing to further analysis or collaboration.

The present version has several limitations. A primary constraint is that the database models are in TypeScript to run on the website itself. This version can either be written manually or, more simply, generated using the MxlPy [13] package, which is available only in Python. While the package is in active development and is already capable of importing an SBML file, this limits the contribution to a Python-favored model construction. The model downloads are also Python-oriented and include a SBML file, which is widely accepted as a model import in most popular programming languages. Another limitation is that validation of the model implementation depends on the information provided in the original publications/provided code; in some cases, exact reproduction may not be possible because simulation settings or experimental inputs are incompletely described. We can only guarantee that our implementation is clearly documented and available. Thus, additional modifications can be implemented quickly if needed. Thirdly, the current focus is on ODE-based mechanistic models for single-cell photosynthesis. It does not yet systematically include constraint-based, agent-based, stochastic, spatial, or empirical modeling frameworks, as well as canopy/ecosystem photosynthesis models. This excludes major approaches such as crop models [23] and global primary production models [24]. The database, however, already includes various versions of the FvCB model [25]. Within the scope of the mechanistic kinetic photosynthesis model, the current database has not been extended to non-C3 photosynthesis in higher plants and to photosynthesis models for other organisms, such as cyanobacteria. These choices were made deliberately to establish a coherent initial scope, but they also define the boundaries of the current resource.

Future development will focus on expanding model coverage, improving metadata completeness, and strengthening validation workflows. Additional functionality may include model comparison tools, standardized export formats, integration with external repositories, and programmatic access through an application programming interface. As the database grows, GreenSloth has the potential to become a central infrastructure for reproducible photosynthesis modeling.

GreenSloth demonstrates that a single curated, standardized resource can simultaneously serve computational biologists seeking reusable and modularized model components, experimental collaborators seeking to test a hypothesis without writing code, and researchers in the scientific machine learning community seeking validated mechanistic building blocks for hybrid modeling. As the database continues to grow through community contribution, we hope it can shift the norm in photosynthesis modeling from isolated, hard-to-reproduce implementations toward shared, actively maintained infrastructure.

## Data availability

The GreenSloth database is freely accessible at https://computational-biology-aachen.github.io/green-sloth/ or https://greensloth.rwth-aachen.de/without registration. The source code, curated model implementations, metadata files, and validation material are available at https://github.com/Computational-Biology-Aachen/green-sloth.

## Author contributions

Elouen Corvest: model curation, software implementation, validation, writing – original draft. Marvin van Aalst: model curation, software implementation, validation, writing – original draft. Anna Matuszyńska: conceptualization, model curation, supervision, funding acquisition, writing – original draft, review and editing. Other authors: implementation and validation of at least two models, writing – review, and editing.

