## Supplemental Table 1 for "GreenSloth: a curated database and executable platform for mechanistic photosynthesis models"

**Table 1:** Content of the GreenSloth database. Models are ordered alphabetically, reflecting the website.

| Model | Organism / system | Process tag(s) | Model type | Data type | Ref. |
|---|---|---|---|---|---|
| Bellasio et al. (2019) | Higher plant / leaf photosynthesis | Gas exchange, carbon assimilation | ODE | Gas exchange | [1] |
| Bernacchi et al. (2013) | $C_3$ plants / photosynthetic carbon assimilation | Carbon assimilation | Steady-state (Farquhar, von Caemmerer, and Berry (FvCB)-based) | Gas exchange | [2] |
| Davis et al. (2017) | Higher plant | *pmf*, bioenergetics | ODE / Steady-state? | No data given | [3] |
| Ebenhöh et al. (2014) | *Chlamydomonas reinhardtii* | Electron transport, NPQ, acclimation | ODE | Fluorescence | [4] |
| Fuente et al. (2024) | Photosynthetic system | Dynamic response, environmental perturbation | ODE | Frequency Domains | [5] |
| Farquhar et al. (1980) | Higher plant / Calvin–Benson cycle | Carbon fixation | Steady-state | Gas exchange | [6] |
| Gu et al. (2023) | Higher plant / Photosynthetic electron transport chain | Linear electron transport, redox reactions | Steady-state | Fluorescence | [7] |
| Hahn et al. (1987) | Higher plant / Calvin–Benson cycle | Carbon fixation, photorespiration | ODE | Concentration changes | [8] |
| Johnson et al. (2021) | Higher plant / leaf photosynthesis | Electron transport, Carbon Fixation | Steady-state | Gas-exchange | [9] |
| Lam et al. (2026) | *Nicotiana benthamiana* | NPQ, photoinhibition | ODE | Fluorescence lifetime | [10] |
| Lazár et al. (1997) | Spring barley (*Hordeum vulgare L. cv. Akcent*) | Fluorescence induction | ODE | Fluorescence (OJIP) | [11] |
| Li et al. (2021) | Photosynthetic system | Photosynthetic regulation | ODE | ECS (P515) / Fluorescence | [12] |
| Matuszyńska et al. (2016) | *Arabidopsis thaliana* | NPQ, photoprotection | ODE | Fluorescence | [13] |
| Matuszyńska et al. (2019) | *Arabidopsis thaliana* | Electron transport, carbon assimilation | ODE | Fluorescence | [14] |
| Morales et al. (2018) | Higher plant / leaf photosynthesis | Carbon assimilation, stomatal conductance, NPQ | ODE | Gas exchange, Fluorescence | [15] |
| Poolman et al. (2000) | Higher plant / Calvin–Benson cycle | Carbon fixation | ODE | Gas exchange / Concentration changes | [16] |
| Saadat et al. (2021) | Higher plant / photosynthetic electron transport | Electron transport, redox regulation | ODE | Fluorescence / P700 absorbance | [17] |
| Salvatori et al. (2022) | Soybean (*Glycine max (L.) Merrill*) | Electron Transport, Assimilation, NPQ | ODE | Gas exchange / Fluorescence | [18] |
| Yokota et al. (1985) | Euglena gracilis | Photorespiration | ODE | Gas exchange | [19] |
| Zaks et al. (2012/2014) | Higher plant / thylakoid membrane | NPQ, *pmf* | ODE | Fluorescence | [20] |
| Zhu et al. (2005) | Higher plant / photosystem II | Electron transport, fluorescence induction | ODE | Fluorescence (OJIP) | [21] |
| Zhu et al. (2009) | Theoretical / Calvin–Benson cycle | Carbon fixation, | ODE | None | [22] |

[10] Lam Lam, Rebecca Lee, Dhruv Patel-Tupper, Henry E. Lam, Tsung-Yen Lee, Alexa Ma, Sophia A. Ma, Hetty He, Krishna K. Niyogi, and Graham R. Fleming. Dissecting the contributions to non-photochemical quenching in a land plant under fluctuating light. *Nature*

*Communications*, 17(1):3664, March 2026. ISSN 2041-1723. doi:10.1038/s41467-026-70414-2. URL https://www.nature.com/articles/s41467-026-70414-2.
